# Causal Effects of Pupil Size on Visual Processing

**DOI:** 10.1101/2023.03.08.531702

**Authors:** Sebastiaan Mathôt, Hermine Berberyan, Philipp Büchel, Veera Ruuskanen, Ana Vilotijević, Wouter Kruijne

## Abstract

The size of the eyes’ pupils determines how much light enters the eye and also how well this light is focused. Through this route, pupil size shapes the earliest stages of visual processing. Yet causal effects of pupil size on vision are poorly understood and rarely studied. Here we report the effects of both experimentally induced and spontaneous changes in pupil size on visual processing as measured through EEG. We compare these to the effects of stimulus intensity and covert visual attention, because previous studies have shown that these factors all have comparable effects on some common measures of early visual processing, such as detection performance and steady-state visual evoked potentials; yet it is still unclear whether these are superficial similarities, or rather whether they reflect similar underlying processes. Using a mix of neural-network decoding, ERP analyses, and time-frequency analyses, we find that induced pupil size, spontaneous pupil size, stimulus intensity, and covert visual attention all affect EEG responses, mainly over occipital and parietal electrodes, but—crucially—that they do so in qualitatively different ways. Induced and spontaneous pupil-size changes mainly modulate activity patterns (but not overall power or intertrial coherence) in the high-frequency beta range; this may reflect a causal effect of pupil size on oculomotor activity and/ or visual processing. In addition, spontaneous (but not induced) pupil size tends to correlate positively with intertrial coherence in the alpha band; this may reflect a non-causal relationship, mediated by arousal. Taken together, our findings suggest that pupil size has qualitatively different effects on visual processing from stimulus intensity and covert visual attention. This shows that pupil size causally affects visual processing, and provides concrete starting points for further study of this important yet understudied earliest stage of visual processing.

---

Visual perception starts as soon as light passes through the eye’s pupil, before it has even reached the retina. By actively controlling the size of the pupil, the visual system controls how much light enters the eye (which increases with increased pupil size), as well as how sharply this light is focused on the retina (which decreases with increased pupil size). In this way, pupil size shapes visual perception already at the earliest stage (Mathôt, 2020). Yet remarkably little is known about *how* pupil size shapes perception—what exactly is different about vision with large pupils as compared to vision with small pupils? The current study is among the first to directly investigate causal effects of pupil size on visual processing as measured through electroencephalography (EEG; also see Bieniek et al., 2013; Bombeke et al., 2016; Suzuki et al., 2019).

Causal effects of pupil size on behavior have already been established in the 1950s, in a series of studies in which pupil size was manipulated experimentally (Campbell, 1957; Campbell & Gregory, 1960; Woodhouse, 1975). For example, in a landmark study, Campbell and Gregory (1960) used eye drops to maximally dilate the pupil, and then placed an artificial pupil (a device with a gap of adjustable size) in front of the eye. Next, for a range of artificial pupil sizes, they established the lowest intensity at which participants were still able to make out the details of stimuli of various sizes, a task that relies both on visual sensitivity (the ability to detect the presence of a stimulus) and acuity (the ability to make out the details of a stimulus). Crucially, the authors found that small stimuli were best perceived with small pupils, whereas large stimuli were best perceived with large pupils. Phrased differently, they found that visual acuity is highest for small pupils, which provide sharper focus, whereas visual sensitivity is highest for large pupils, which provide increased light influx. This general pattern has since received strong support from fundamental vision science (Eberhardt et al., 2022; Liang & Williams, 1997; Mathôt & Ivanov, 2019), as well as from research on ergonomics of display design (Piepenbrock et al., 2014) and ophthalmology (Alfonso et al., 2007).

Causal effects of pupil size on EEG measures have only been investigated in a handful^1^ of studies (Bieniek et al., 2013; Bombeke et al., 2016; Suzuki et al., 2019). The most compelling example to date is perhaps a study by Suzuki, Minami, and Nakauchi (2019), who recorded steady state visual evoked potentials (SSVEPs) triggered by rhythmically flashing dots; they manipulated pupil size by presenting these dots either on top of a control stimulus or a “glare stimulus”, which is an optical brightness illusion that triggers pupil constriction (Laeng & Endestad, 2012). SSVEPs correspond to rhythmic EEG activity at the frequency of the triggering stimulus, with higher oscillatory power indicating stronger visual responses (Müller et al., 1998). Crucially, the authors found that larger pupils resulted in stronger SSVEPs.

The studies reviewed above suggest that the size of the pupil influences the strength of incoming visual stimuli. Superficially then, the effect of increased pupil size seems similar to the effect of increased stimulus intensity as well as the effect of covert visual attention: at the behavioral level, these two factors also increase the detectability of stimuli (Carrasco et al., 2000); and at the neural level, these two factors also increase SSVEP amplitude (Andersen et al., 2012). This simple observation inspired us to directly compare the effects of pupil size to the effects of stimulus intensity and covert visual attention on visual processing, as a crucial first step towards understanding the role of pupil size in visual processing.

We used a spatial cueing paradigm in which we manipulated covert visual attention by presenting the target either at a cued (attended) or uncued (unattended) location. We manipulated stimulus intensity by having the target be either bright or dim. And we manipulated pupil size using a novel technique from our lab that relies on isoluminant red and blue inducers, while avoiding any spatial or temporal overlap between the inducers and the other stimuli (as described in more detail under <u>Pupil-size induction procedure</u>); for reference, this technique induces substantial differences in pupil size (about 20% surface area) that are far larger than the effects of arousal or mental effort, yet smaller than the effect of direct exposure to light. We further included spontaneous fluctuations in pupil size as a pseudo-manipulation to allow for better comparison to previous studies that looked at correlations between spontaneous pupil-size fluctuations and EEG measures (e.g., Hong et al., 2014; Murphy et al., 2011; Podvalny et al., 2021; Thigpen et al., 2018; Waschke et al., 2019; for related non-human animal studies, see Franke et al., 2022; Joshi et al., 2016).

We will focus on EEG activity after target presentation, as opposed to pre-target or baseline activity, and on main effects of induced pupil size, spontaneous pupil size, stimulus intensity, and covert attention, as opposed to interactions between these factors. Our data set, which is publicly available and well-documented for re-use (see <u>Open-practices statement</u>), allows for many more analyses; however, because there is limited previous research to build on, we believe that this clear focus will better allow us to draw conclusions and to formulate concrete hypotheses for further investigations.

## Methods

### Participants

Thirty participants (26 females; M*_age_* = 20.73) were included in the experiment, which was approved by the local ethics board of the University of Groningen, Netherlands (approval code: PSY-2122-S-0341). All participants had normal or corrected-to-normal vision, were right-handed, and had no history of neurological disorders. Participants received course credit and provided written informed consent. One additional participant was tested but excluded due to a technical recording error. Two additional participants were tested but excluded because they did not show a pupil-size-induction effect (see <u>Pupil-size induction procedure</u>). These three participants were replaced to reach our predetermined sample size of thirty participants. In the absence of an a-priori expected effect size, our sample size was based on the rule of thumb provided by Brysbaert and Stevens (2018) to have at least 1,600 observations per condition; in our design, we had 5,760 observations per condition when considering main effects.

### Materials and software

The experimental script was implemented using OpenSesame 3.3 (Mathôt et al., 2012) using the PsychoPy (Peirce, 2007) backend for display presentation and PyGaze (Dalmaijer et al., 2014) for eye tracking.

The experiment was conducted on a desktop computer with a 27” flatscreen monitor with a refresh rate of 60 Hz and a resolution of 1920 × 1080 pixels. The right eye was recorded at 1000 Hz using an EyeLink 1000 (SR Research). EEG data was recorded at 1000 Hz using a TMSI REFA 32 system with 26 channels placed according to the standard 10-20 system with OpenViBE data acquisition software (Renard et al., 2010). Two additional channels were placed on the left and right mastoids for offline re-referencing. Two additional channels were placed above and below the left eye; the difference between these channels was determined offline to serve as a vertical EOG channel. Two additional channels were placed on the left and right temples; their difference served as a horizontal EOG channel (see also <u>Data preprocessing</u>).

Data analysis was performed using the Python `eeg_eyetracking_parser`^2^ toolbox, which is a high-level toolbox that uses `MNE` (Gramfort et al., 2013) for general EEG processing, `eyelinkparser` (Mathôt & Vilotijević, 2022) for eye-movement and pupil-size processing, `autoreject` (Jas et al., 2017) for rejecting and repairing bad EEG channels and epochs, and `braindecode` (Schirrmeister et al., 2017) for neural-network based decoding. Bayesian analyses were performed using JASP (JASP development team, 2021). Cluster-based permutation tests and cross-validation tests were performed using `time_series_test` (Mathôt & Vilotijević, 2022).

### Pupil-size induction procedure

We manipulated pupil size by presenting red and blue inducers before each trial, where we expected pupils to be larger after sustained exposure to red inducers as compared to blue inducers. Crucially, this pupil-size difference should not result from differences in luminance between red and blue inducers, which would affect the initial pupil response to a brief light stimulus (< 2s), but rather from differential activation of intrinsically photosensitive retinal ganglion cells (ipRGCs), which affects the long-term pupil response to a sustained light stimulus (> 10 s; Gamlin et al., 2007; Mathôt, 2018). Therefore, we obtained equiluminant intensities of red and blue for each participant separately during a luminance-calibration phase at the start of the experiment (see also Kinzuka et al., 2022; Wardhani et al., 2022). The aim of this calibration phase was to find intensities of red and blue that would result in an equally strong initial pupil constriction and thus be equiluminant in this specific sense.

During luminance calibration, participants looked passively at a central fixation cross while they were exposed to peripheral red or blue stimuli that were presented for 1000 ms followed by 2000 ms of a dark (0.10 cd/m^2^) screen; the center (*r* = 8.36°) of the display remained dark at all times (Fig. 1a). Red and blue stimuli were presented in alternation. The intensity of the blue stimulus was fixed (9.84 cd/m^2^). The intensity of the red stimulus was adjusted online based on the strength of pupil constriction in response to the red or blue stimulus; specifically, on each trial, the strength of pupil constriction was calculated as the difference between the maximum and minimum pupil size between 900 and 1,800 ms after the onset of the stimulus. If pupil constriction was weaker in response to a red stimulus than in response to the preceding blue stimulus, or stronger in response to a blue stimulus than in response to the preceding red stimulus, then the intensity of the red stimulus was increased by 1%; otherwise, the intensity of the red stimulus was decreased.

**Figure 1.**
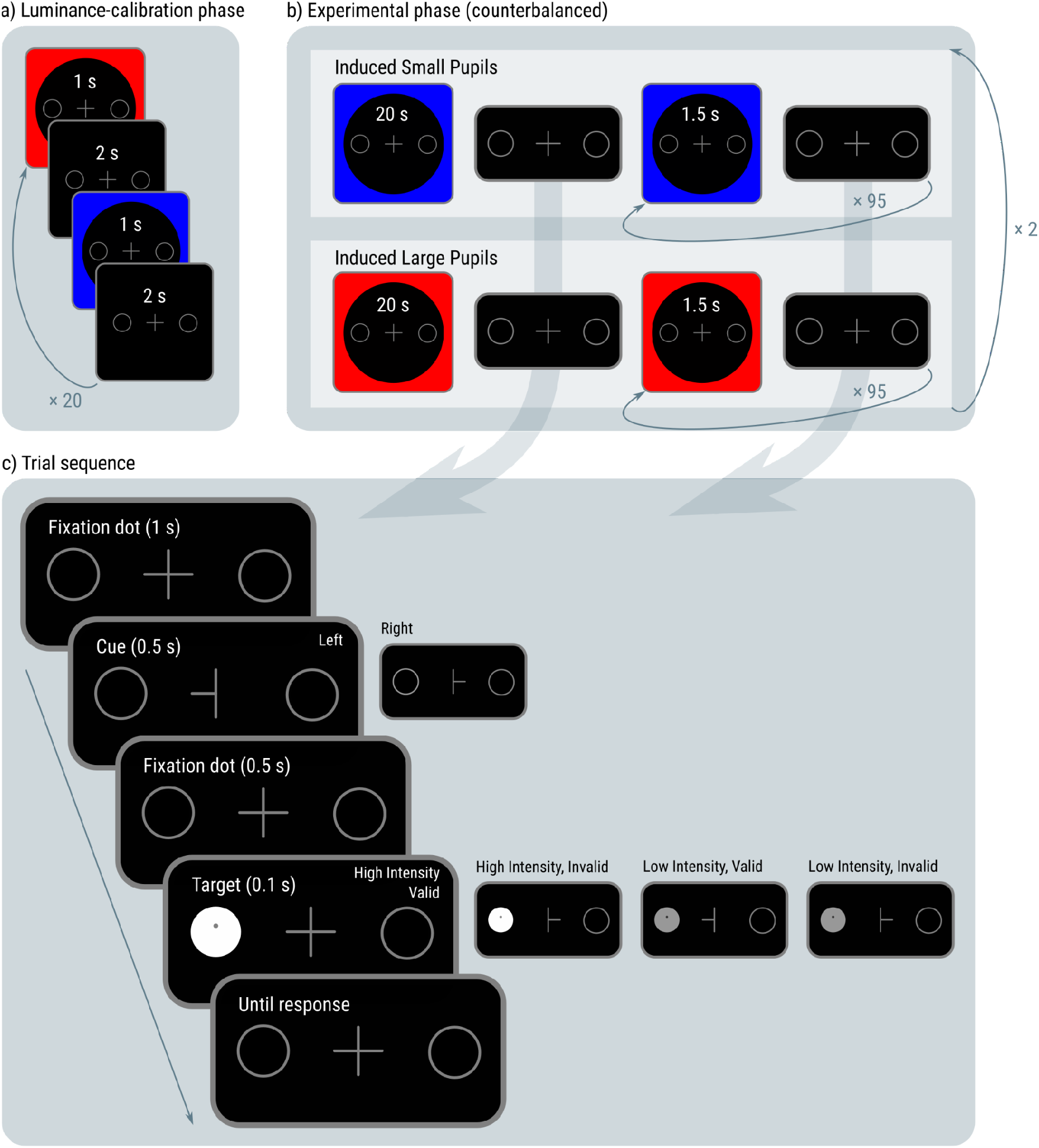
a) During luminance calibration, participants passively viewed alternating red and blue displays. Red intensity was adjusted until red and blue displays triggered equally strong pupil constriction. b) The experiment consisted of two red-inducer (large pupils) blocks and two blue-inducer (small pupils) blocks. The first inducer of each block was longer (20 s) than subsequent inducers (1.5 s). c) During each trial, participants reported whether a small dot (the target) was slightly above or below the midline. Target intensity and cue validity were manipulated.

The final participant-specific red (adjusted) and blue (fixed) intensities were used for the inducer stimuli during the experiment (Fig. 1b). Specifically, the first trial of each block of 96 trials started with a red or blue inducer that was presented for 20 s, which is long enough to trigger a substantial difference in the sustained pupil response such that pupils are larger after red inducers than after blue inducers (Mathôt, 2018; Wardhani et al., 2022). Subsequent trials in the same block started with a ‘top-up inducer’ of the same color that was presented for 1.5 s plus the duration of the drift-correction procedure. Such a brief presentation time is by itself not sufficient to trigger a difference in pupil size between red and blue inducers; however, we found in pilot studies (data not included) in which we tried various induction procedures on ourselves that a brief top-up inducer is able to partly reinstate the effect of a previously presented long inducer. We relied on this effect to reduce the overall duration of the experiment.

Each participant completed two practice blocks of 10 trials and four non-practice blocks of 96 trials, during which the inducer color was kept constant. The block order was either red-blue-red-blue or blue-red-blue-red, counterbalanced across participants.

### Experimental procedure

Following the inducer, each trial started with a dark (0.10 cd/m^2^) display consisting of a central fixation dot and two circular placeholders (*r* = 1.49°), one on the left and one on the right side of the display (eccentricity = 5.02°), presented for 1000 ms (Fig. 1c). Except for the target (see below), stimuli were low intensity (0.66 cd/m^2^). Next, a cue was presented for 500 ms; the cue was identical to the fixation cross, except that either the left or right arm disappeared, indicating that the target was most likely (75%) to appear at the side of the remaining arm. Next, the fixation cross re-appeared for 500 ms, after which the target was presented for 100 ms while the cue remained on the display; the target was a small dark dot that appeared slightly above or below the vertical midline of the screen, inside a bright (83.58 cd/m^2^) or dim (7.96 cd/m^2^) filled circle that appeared at the location of one of the placeholders. Next, the fixation cross and placeholders re-appeared until the participant responded or until a timeout of 2,000 ms occurred. The participant’s task was to report whether the target dot appeared above or below the midline by pressing the ‘m’ or ‘n’ key as fast and accurately as possible. Participants were instructed to keep their eyes fixated on the central fixation cross throughout the trial.

Throughout the experiment, a continuous 2-up-1-down staircase ensured that performance for valid trials remained around 66% by making the target appear closer to (more difficult) or further away from (easier) the midline. Initial staircase values were set to a medium level of difficulty (0.5), and converged to a participant-specific difficulty level during the practice phase. The staircase was run independently for red and blue blocks to ensure that task difficulty did not differ between red and blue blocks.

The experimental design consisted of three experimental factors: induced pupil size (large, small; varied between blocks), stimulus intensity (bright, dim; varied within blocks), and covert visual attention (attended, unattended; varied within blocks). We also treated spontaneous pupil size (large, small) as a pseudo-experimental factor. This factor encoded whether the mean pupil size during the presentation of the first fixation display of each trial was larger or smaller than the per-participant, per-inducer-color median pupil size; in other words, this factor encoded spontaneous fluctuations in pupil size in a way that was uncorrelated with induced changes in pupil size. Together, these factors form a 2 (induced pupil size) × 2 (spontaneous pupil size) × 2 (stimulus intensity) × 2 (covert visual attention) within-subject design.

### Data preprocessing

#### EEG preprocessing

EEG data was preprocessed fully automatically. Unless otherwise specified, we used the default parameters as described on the documentation of the referenced functions. 1) Data was re-referenced to the mastoid channels. 2) Muscle artifacts, which are characterized by bursts of high-frequency activity (110 - 140 Hz), were marked as bad using the MNE function for annotating muscle activity (see `mne.preprocessing.annotate_muscle_zscore`) with a z-threshold of 5. 3) Data was downsampled to 250 Hz for computational efficiency. 4) A vertical EOG channel was created by subtracting the electrodes above and below the left eye, and a horizontal EOG channel was created by subtracting the electrodes on the left and right temples. 5) Data was filtered using a 0.1 - 40 Hz bandpass filter. 6) The RANSAC algorithm (see `autoreject.Ransac`) determined bad channels based only on data segments corresponding to trials; in brief, this algorithm assumes that a channel is bad if its data is typically poorly predicted by interpolation from neighboring channels (Bigdely-Shamlo et al., 2015). Overall, 2.6% of channels were marked as bad. 7) The influence of blinks and eye movements was reduced by first running an independent component analysis (ICA) on a 1 Hz highpass filtered copy of the data, then identifying those ICA components that correlated highly with the EOG channels, and finally removing these components from the data (see `mne.preprocessing.ICA`). Overall, 7.2% of ICA components were removed. 8) Channels that were marked as bad in step 6 were interpolated (see `mne.io.Raw.interpolate_bads`). 9) Epochs containing saccadic eye movements larger than 3.0° as identified by the eye tracker were marked as bad and excluded from analysis. Overall, 14.7% of epochs were marked as bad based on the presence of saccadic eye movements and/ or muscle activity (as described above).

#### Additional preprocessing for EEG decoding

For the decoding analyses, we followed the recommendations from Schirrmeister et al. (2017) and the associated `braindecode` documentation^3^. Specifically, a 4 - 30 Hz bandpass filter was applied to reduce artifacts, and data was transformed using exponential moving standardization (see `braindecode.preprocessing.exponential_moving_standardize`). Data was augmented using cropped decoding with a window of 800 ms (within a 850 ms epoch) and a window stride of 4 ms; that is, overlapping 800 ms windows were cut from each epoch (0 - 800 ms, 4 - 804 ms, etc.) and these windows were used as separate pieces of training data. Cropped decoding is a way to artificially increase the amount of training data (“data augmentation”), which is common practice in training of neural networks. Two participants were excluded from the decoding analyses, because they missed valid trials for one of the sixteen combinations of factors. (This was not an issue for the non-decoding analyses, which were all trial-level analyses that are robust to missing data.) For the other participants, when conditions did not have an equal number of trials, trials were randomly sampled with replacement from the least frequent condition until the number of trials was equal for all conditions during training (but not for testing, which does not require balanced data).

#### Additional preprocessing for ERP and time-frequency analyses

For the ERP and time-frequency analyses, the Autoreject algorithm (see `autoreject.Autoreject`) was applied to the remaining data to detect and interpolate epochs and channels that had not been identified in the previous preprocessing steps. ERPs were baseline corrected relative to a 100 ms pre-target interval. Importantly, we did *not* apply baseline correction for the time-frequency analyses. This choice was based on the fact that induced pupil size was constant within blocks (see Fig. 2b), and could thus affect frequency power and inter-trial coherence already before target onset; because of this, baseline correction could make pre-target (baseline) differences between large and small induced pupils appear as post-target (evoked) differences, which could lead to incorrect interpretations.

**Figure 2.**
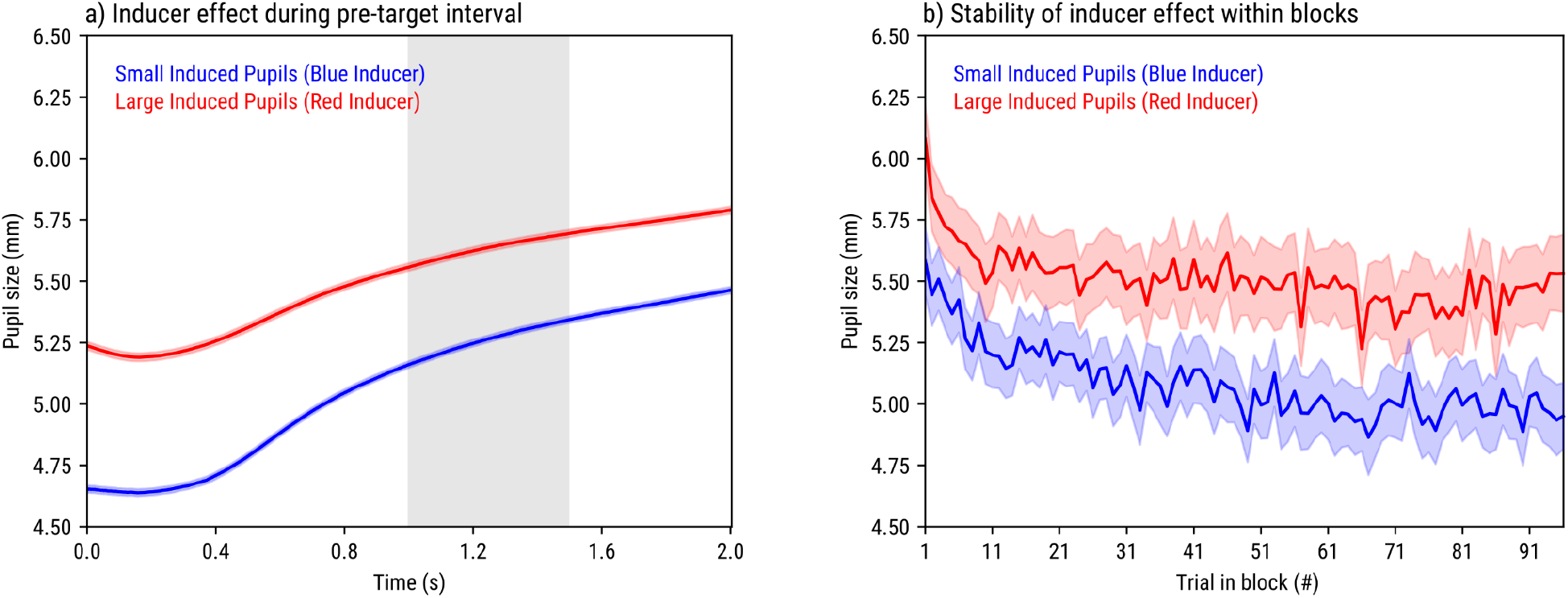
The effect of red / blue inducers on pupil size. Error bands indicate grand standard error. a) From the start of the trial until the presentation of the target. The gray band indicates the presentation of the cue. b) Mean pupil size during the pre-target interval as a function of trial position in block and inducer color.

#### Pupil-size preprocessing

For the pupil-size analyses, we followed the recommendations from Mathôt and Vilotijević (2022). 1) Missing or invalid data was interpolated using cubic-spline interpolation if possible, using linear interpolation if cubic-spline interpolation was not possible (when the segment of missing data was too close to the start or end of a trial), and removed if interpolation was impossible altogether (when data was missing from the start and/ or until the end of a trial) or if the period of missing data was longer than 500 ms and thus unlikely to reflect a blink (see `datamatrix.series.blinkreconstruct`). 2) Pupil size was converted from arbitrary units as recorded by the EyeLink to millimeters of diameter. 3) Epochs for which baseline pupil size (the average of the 50 ms before target onset) was missing due recording artifacts or deviated more than 2 SD from the mean baseline pupil size of that participant were excluded from analysis. Overall, 17.2% of trials were marked as bad based on this criterion, roughly equally divided across most factors, with the exception that trials with spontaneously large pupils (22.6%) were more often marked as bad than trials with spontaneously small pupils were (11.9%); this likely reflects that participants were more likely to move (and thus distort the recording) when they were in a state of increased arousal.

#### Reporting of statistical analyses

We will report a large number of statistical analyses, many of which are fully or partly exploratory, and none of which were pre-registered. We will therefore emphasize general patterns over significance of individual effects. When reporting statistical analyses, we will refer to *p* < .05 as ‘significant effects’ (i.e., using an alpha level of .05). We will focus on main effects.

## Results

### Red and blue inducers successfully manipulated sustained pupil size

Pupils were larger in red-inducer blocks as compared to blue-inducer blocks (see Fig. 2a); that is, even though the strength of the *initial* pupil constriction to both inducer colors had been matched by our luminance-calibration procedure (see <u>Supplementary results</u>), *sustained* pupil size differed systematically. This was true for all participants that were included in the analysis.^4^ The inducer effect was stable across blocks (see Fig. 2b), suggesting that the brief top-up inducers that preceded each trial were sufficient to reinstate the effect of the 20 s inducer that preceded each block. Since we had selected participants based on whether or not they showed this effect no statistics were performed on this result.

### EEG: decoding

As a first step, we used neural-network decoding to investigate whether and how the ERP signal around target onset was affected by our four factors: induced pupil size, spontaneous pupil size, stimulus intensity, and covert visual attention. We focused on the signal from 100 ms before until 750 ms after target onset (see also <u>Data preprocessing</u>). All decoding analyses were done using the ‘shallow’ neural network from the `braindecode` toolkit using the network’s default parameters (Schirrmeister et al., 2017). This is a four-layer network that first performs a temporal filter for each channel separately, followed by a spatial filter that combines information across channels, thus allowing the network to identify both temporal and spatial regularities in the signal. We chose this ‘shallow’ network over the toolkit’s ten-layer ‘deep’ network to reduce training time. To assess the generalizability of the decoding, we used four-fold cross-validation for each participant separately.

In a series of analyses, we tested for each factor: a) whether it could be decoded at all; b) which electrodes contributed most to decoding; and c) which frequencies contributed most. Since neural network-based decoding requires fairly long epochs as input, as opposed to single samples as is common for other EEG-decoding techniques (see e.g., Grootswagers et al., 2017 for an overview), we did not attempt to test which time points within this window contributed most. Finally, we tested d) whether any pair of factors are similar by training on one factor and then decoding on another (cross-decoding).

#### All factors can be reliably decoded (Fig. 3)

We first performed overall decoding of all 16 combinations of factors. A confusion matrix revealed that all factors were reliably decoded (visible as the red diagonal in Fig. 3a) with an average accuracy of 42.90% (Fig. 3b), far above the 6.25% chance level (one sample t-test against chance level: *t* = 5.68, *p* < .001). Decoding accuracy for an individual factor was derived from the full confusion matrix by averaging across all other factors, resulting in a 2×2 confusion matrix for each factor; this showed that individual factors were similarly decoded far above the 50% chance level: induced pupil size (77.52%, *t* = 16.22, *p* < .001), spontaneous pupil size (70.54%, *t* = 18.09, *p* < .001), target intensity (69.14%, *t* = 21.06, *p* < .001), and covert visual attention (86.48%, *t* = 59.30, *p* < .001).

**Figure 3.**
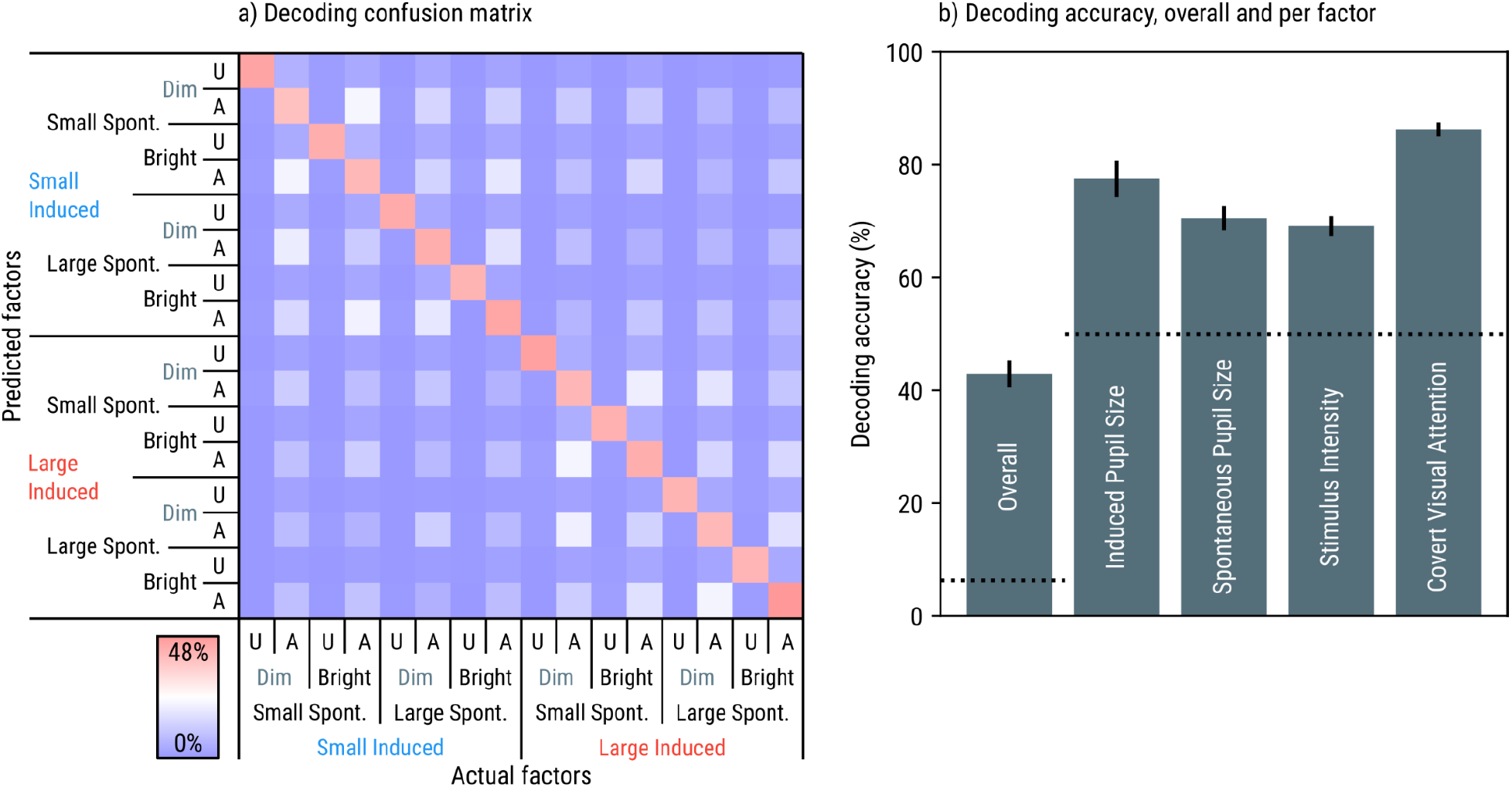
Main results of decoding target-evoked responses. a) Confusion matrix with actual factors on the x-axis and predicted factors on the y-axis. U = Unattended, A = Attended, Spont. = Spontaneous. High (red) values correspond to frequently predicted factors given an actual factor; low (blue) values correspond to infrequent predictions. Percentages sum to 100% per column. The pronounced red diagonal reflects that factors were reliably decoded. b) Decoding accuracy overall and per factor. The dotted horizontal lines indicate chance level. Error bars indicate 95% confidence intervals.

Because pupil-size inducers were varied between blocks, decoding of induced pupil size was confounded by proximity in time: two trials with the same inducer color were on average closer to each other in time than two trials with different inducer colors were. The Supplementary Results contain several control analyses showing that induced pupil size can be reliably decoded also in decoding schemes in which time is no longer a confounding factor (see <u>Decoding of</u> <u>induced pupil size does not rely on proximity in time</u>).

#### Occipital and parietal channels carry most information for all factors (Fig. 4)

To identify which electrodes contributed most to decoding (i.e., were most strongly affected by our factors), we conducted an ICA-based perturbation analysis for each participant and factor separately; this analysis is documented in detail in the analysis source code (see <u>Open-practices statement</u>), but in brief, we determined how much decoding performance dropped after removing a single independent component from the signal for both training and testing. The rationale behind this procedure is that by removing independent components rather than electrodes from the signal, this ICA-based perturbation analysis is less hindered by high correlations between neighboring electrodes. We then assigned a ‘contribution score’ to each electrode by multiplying this drop in decoding accuracy by the (absolute) loading of the electrode onto the excluded independent component, and dividing the result by the original decoding accuracy. We repeated this procedure for each of the 26 independent components, each time increasing the contribution scores for the electrodes. In the end, electrodes that carried most information thus accumulated the highest contribution scores. (The analysis source code contains a sanity-check analysis showing that applying this ICA-based perturbation analysis to a dummy factor that is randomly assigned to trials indeed results in a flat scalp topography.)

**Figure 4.**
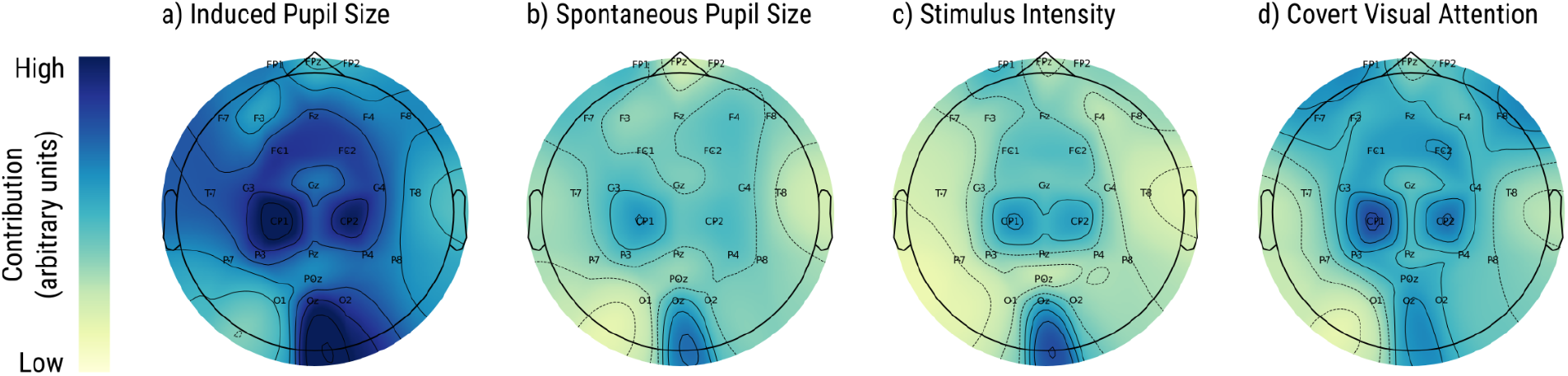
Contributions of electrodes to decoding of a) induced differences in baseline pupil size, b) spontaneous differences in baseline pupil size, c) stimulus intensity, and d) covert visual attention. Dark blue values indicate strong contributions; green-yellow values indicate weak contributions.

We subsequently conducted a repeated measures ANOVA using the contribution score as the dependent variable, and factor and electrode as independent variables; to qualify the results, we relied on visual inspection of scalp topographies (Fig. 4). We found a main effect of electrode (F(25, 675) = 3.87, *p* < .001), reflecting that occipital and parietal electrodes contributed most to decoding overall, with CP1 and CP2 having the highest contribution scores. We did not find a reliable main effect of factor (F(3, 81) = 1.77, *p* = .160), reflecting that, on average, removing independent components from the signal impaired decoding of each factor about equally. We also did not find a reliable electrode × factor interaction (F(75, 2025) = 1.24, *p* = .080), reflecting that there was no notable difference between the four factors in which electrodes contributed most to decoding.

#### Induced and spontaneous differences in pupil size mainly affect beta; stimulus intensity mainly affects alpha; covert visual attention mainly affects theta (Fig. 5)

To identify which frequencies carried most information, we conducted a frequency-based perturbation analysis for each participant and factor separately; this analysis was similar to the ICA-based perturbation analysis, except that we now assigned a contribution score to specific frequencies, based on how much decoding performance was impaired after silencing these frequencies using a notch filter. We used a notch-filter that increased exponentially in 15 steps from 4 (lower theta) to 30 (upper beta) Hz, using a filter width that similarly increased exponentially from 1 to 7.5 Hz. This frequency range was determined by the bandpass filter that was applied during preprocessing for decoding (see <u>Data preprocessing</u>). (The analysis source code contains a sanity-check analysis showing that applying this frequency-based perturbation analysis to a dummy factor that is randomly assigned to trials indeed results in a flat distribution where no frequency contributes more than others.)

**Figure 5.**
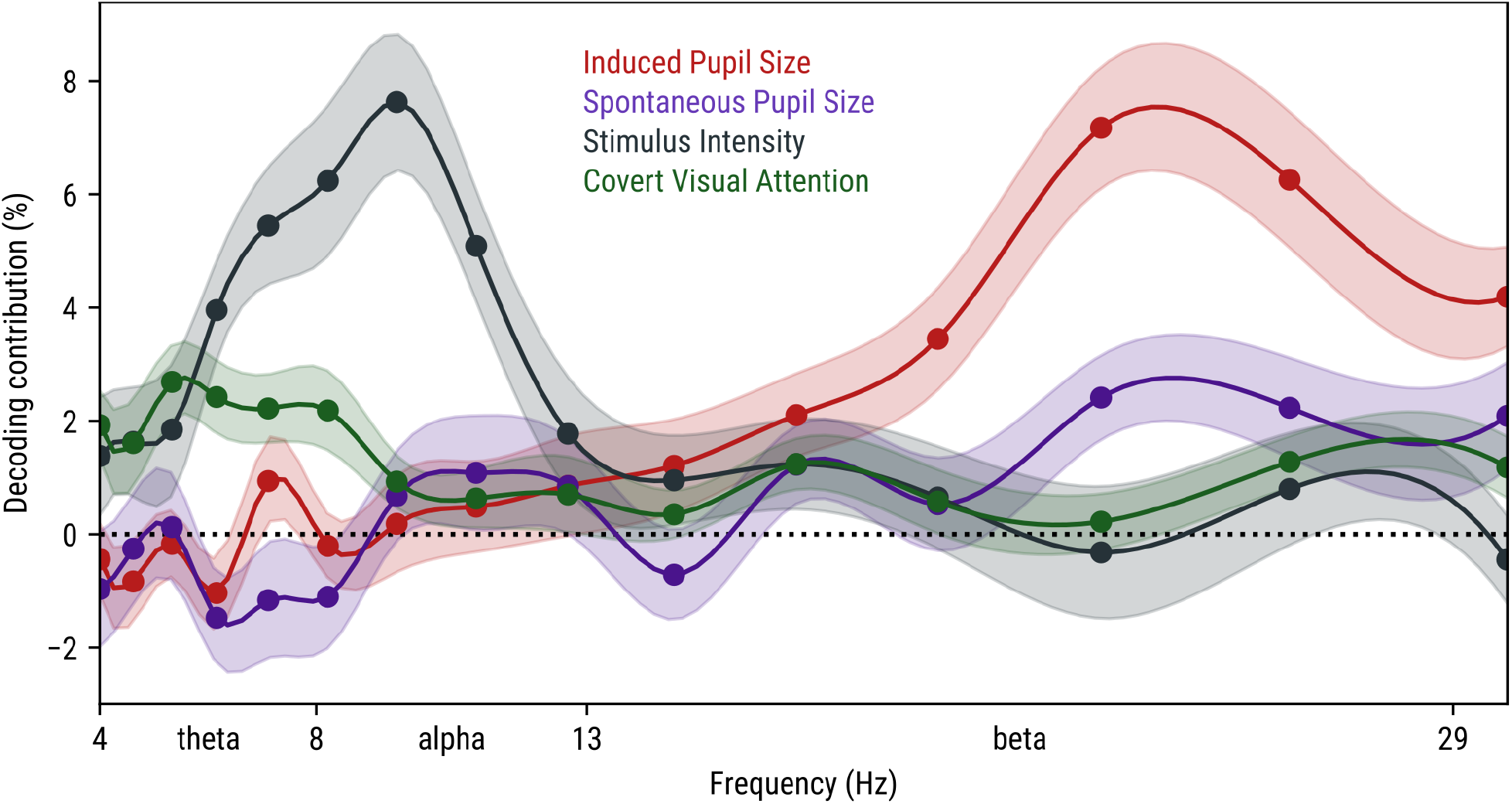
Contribution of frequency bands to decoding of: induced differences in baseline pupil size (red), spontaneous differences in baseline pupil size (purple), stimulus intensity (gray), and covert visual attention (green). The x-axis indicates frequencies. The y-axis indicates contribution scores, which reflect how much decoding accuracy decreased after notch-filtering specific frequencies. Error bands indicate within-subject standard errors. Lines have been smoothed using cubic-spline smoothing.

Two important caveats to guide the interpretation of these results: 1) If silencing a specific frequency strongly impairs decoding, this can reflect several things, including overall power or intertrial-coherence differences as in a traditional time-frequency analysis (David et al., 2006), but also modulation of specific ERPs that are most pronounced in this frequency range; for example, an effect on the P3 component, which spans a relatively long time window, might express itself as a reliance on low frequencies, whereas an effect on the N1 component, which spans a shorter time window, might express itself as a reliance on high frequencies. 2) The decoding-contribution value is a relative within-factor measure, such that differences between frequencies for a given factor are meaningful, but differences between factors for a given frequency are *not* meaningful.

We subsequently conducted a repeated measures ANOVA using the contribution score as a dependent variable, and factor and frequency as independent variables; to qualify the results, we relied on visual inspection (Fig. 5) and simple-effect analyses. We found a main effect of frequency (F(14, 378) = 4.23, *p* < .001), reflecting that some frequencies were more important than others for decoding overall. We did not find a reliable main effect of factor (F(3, 81) = 2.72, *p* = .051), reflecting that there was no notable difference in how much decoding of the four factors was disrupted overall in the analysis (as for the ICA-perturbation analysis). Crucially, we found a factor × frequency interaction (F(42, 1134) = 9.25, *p* < .001), reflecting that there was a difference between the four factors in which frequencies were most important for decoding.

We qualified the interaction with follow-up simple effects analyses. This revealed effects of frequency on induced pupil size (F(14, 378) = 14.45, *p* < .001), which was mainly reliant on beta frequencies with a peak around 22 Hz; on spontaneous pupil size (F(14, 378) = 3.46, *p* < .001), which was also mainly reliant on beta frequencies with a peak around 22 Hz, but less strongly, and with a second, smaller reliance on the alpha band; on stimulus intensity, (F(14, 378) = 8.00, *p* < .001), which was mainly reliant on theta and alpha frequencies with a peak around 10 Hz; and on covert visual attention (F(14, 378) = 3.84, *p* < .001), which was mainly reliant on theta frequencies with a peak around 5 Hz.

#### Cross-decoding reveals qualitatively different effects of all factors

We conducted a cross-decoding analysis by training the neural network to decode one factor (e.g., induced pupil size), using the same training procedure as before, and then using this pre-trained neural network to decode another factor (e.g., stimulus intensity). For each combination of factors, we conducted this analysis in both directions (e.g., training on induced pupil size and testing on stimulus intensity, and then the other way around) and took the average accuracy as our measure of cross-decoding accuracy, where deviations from chance in either direction would indicate cross-decoding.

We subsequently conducted a one-sample *t*-test against chance level (50%) for each combination of factors. This did not reveal reliable cross-decoding for any combination of factors^5^: induced pupil size ↔ spontaneous pupil size (M = 49.9%, *t* = −0.36, *p* = .721); induced pupil size ↔ stimulus intensity (M = 49.9%, *t* = −0.32, *p* = .754); induced pupil size ↔ covert visual attention (M = 49.5%, *t* = −1.14, *p* = .266); spontaneous pupil size ↔ stimulus intensity (M = 49.2, *t* = −1.33, *p* = .196); spontaneous pupil size ↔ covert visual attention (M = 48.9%, *t* = −1.54, *p* = .135); stimulus intensity ↔ covert visual attention (M = 50.1%, *t* = 0.48, *p* = .635).

#### Summary of decoding results

We found that induced changes in pupil size, spontaneous changes in pupil size, stimulus intensity, and covert visual attention could all be reliably decoded from the EEG signal following target onset. In terms of scalp topography, occipital and parietal electrodes contributed most to decoding for all factors. In terms of frequencies, beta frequencies contributed most to decoding of induced and, to a lesser extent, spontaneous changes in pupil size; alpha frequencies contributed most to decoding of stimulus intensity, and again to a lesser extent, spontaneous changes in pupil size; and theta frequencies contributed most to decoding of covert visual attention. There was no cross-decoding between any combination of factors, suggesting that all factors had qualitatively different effects on the EEG signal following target onset.

### EEG: lateralized ERPs

In the previous section, we focused on decoding analyses, which are a powerful tool to detect whether information is present, and (if it is) in which electrodes and frequencies the information is most prominent. However, decoding results can be difficult to interpret and relate to previous EEG studies, many of which have used more traditional analyses. Therefore, we also conducted ERP and (further down) time-frequency analyses in order to further characterize the effects of pupil size on visual processing.

For the ERP analyses, we focused on lateralized (non-midline) parietal electrodes (P3, P4, P7, P8, CP1, and CP2), because our decoding analysis had revealed parietal electrodes, especially CP1 and CP2, as being strongly affected by our factors. We decided to exclude occipital electrodes from this analysis because these are characterized by qualitatively different waveforms than the parietal electrodes. However, analyses of different electrode groups can be found in the analysis source code. In addition, we focused on lateralized ERPs, that is, the difference between electrodes that were contra- and ipsilateral to the target (contra - ipsi), since we had an unbalanced display in which the target was presented only on one side of the display. We analyzed the signal from 100 ms before to 500 ms after target onset; this is a slightly shorter epoch than we had used for the decoding analysis, a choice that was based on the assumption that ERP effects were most likely to arise early in time, and on the need to use a restricted the time window in order to maximize statistical sensitivity.

We conducted cluster-based permutation tests on the lateralized ERPs (see `time_series_test.lmer_permutation_test`). Because we used single-trial linear mixed-effects analyses (which are time-consuming) as the basis for the test, a unified cluster-based permutation test with all factors and their interactions as fixed effects and a maximum random-effects structure would have been prohibitively computationally intensive; we therefore conducted four separate tests, one for each factor as fixed effect. By-participant random intercepts and slopes were included. Clusters were identified using a *p* < .05 criterion, and the absolute sum of z-scores was used as test statistic. We used windows of 16 ms and ran 1,000 iterations for each test. Note that the timing of significant clusters provides only a very rough indicator of when effects arise (Sassenhagen & Draschkow, 2019).

#### Induced differences in pupil size affect ERPs (Fig. 6a)

ERPs were slightly less lateralized for induced large pupils as compared to induced small pupils. The main cluster was around 350 ms (*p* = .039). Notably, there was a period between roughly 250 - 400 ms that was characterized by fluctuations of around 20 Hz (beta band) and that included this main cluster; this aligns with the decoding results, which pointed towards beta frequencies as carrying most of the information about induced pupil size.

**Figure 6.**
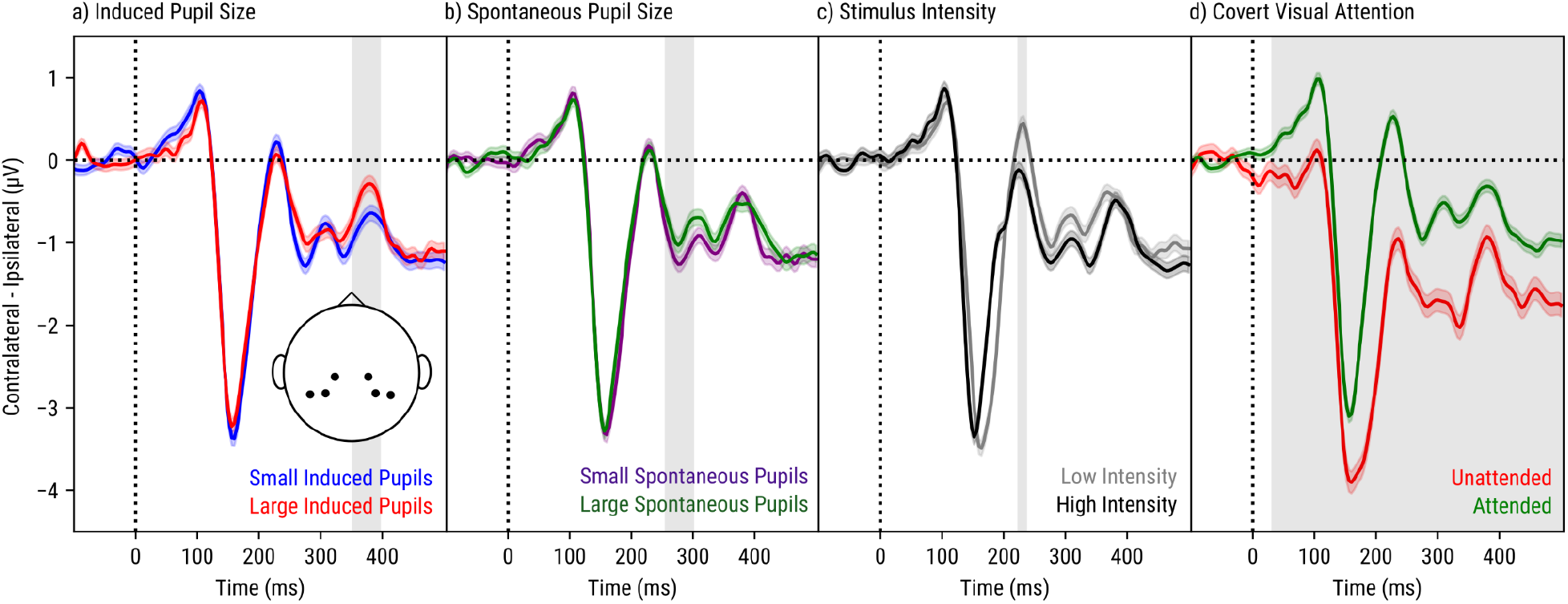
Lateralized target-evoked ERPs (baseline corrected) as a function of time (x-axis) and a) induced pupil size, b) spontaneous pupil size, c) stimulus intensity, and d) covert visual attention. Error bands indicate grand standard errors. Vertical bands indicate significant clusters of differences (see main text for details).

#### Spontaneous differences in pupil size affect ERPs (Fig. 6b)

ERPs were also slightly less lateralized for spontaneous large pupils as compared to spontaneous small pupils. This effect was qualitatively similar to that of induced pupil size, although the main cluster was slightly earlier, around 280 ms (*p* = .044), though still within the period that was characterized by beta oscillations.

#### Stimulus intensity affects ERPs (Fig. 6c)

Stimulus intensity had a complex effect on lateralized ERPs, which is best described as a decreased latency and increased lateralization for increased intensity. The main cluster was around 250 ms (*p* < .001). Of note, the effect of stimulus intensity was qualitatively different from that of induced and spontaneous pupil size.

#### Covert visual attention affects ERPs (Fig. 6d)

By far the most pronounced effect was found for covert visual attention, such that ERPs were less lateralized in response to attended as compared to unattended targets. The main cluster arose very rapidly after target onset and persisted until the end of the signal (*p* < .001). Of note, the effect of covert visual attention was also qualitatively different from that of induced and spontaneous pupil size.

### EEG: time-frequency analyses

For the time-frequency analyses, we again focused on parietal electrodes (P3, P4, P7, P8, CP1, CP2, Pz, and POz), this time also including midline electrodes because we did not focus on lateralized responses (unlike for the ERP analyses)^6^. The analyses were based on Morlet wavelets of two cycles. Frequencies ranged from 4 to 30 Hz in 1 Hz steps. Power was z-transformed for each participant and frequency separately, such that the grand mean across trials, samples, and electrodes was 0 with a standard deviation of 1. We additionally created aggregate time series for power in the theta (4 - 8 Hz), alpha (8 - 13 Hz), and beta (13 - 29 Hz) frequency bands by averaging over subsets of the 1 Hz frequency bands from the Morlet wavelets, and conducted permutation tests on each band separately as described above for the lateralized ERPs. We also extracted intertrial-coherence effects for each band, participant, and factor separately; these were again tested with cluster-based permutation tests, although this time using standard regression analyses (as opposed to linear mixed-effects analyses) since intertrial coherence is an aggregate measure (as opposed to a single-trial measure). As for the ERP analyses and for the same reasons, we analyzed the window from 100 ms before to 500 ms after target onset.

We did not apply baseline correction, nor did we subtract the mean evoked response from the signal; consequently, the analyses reflect both induced (sustained patterns that are not clearly time-locked) and evoked (transient patterns that are locked to stimulus onset) activity (David et al., 2006).

#### Induced differences in pupil size do not reliably affect power or coherence in any frequency band (Fig. 7a,e)

Although the decoding results suggested that beta frequencies, mainly around 22 Hz, were strongly affected by induced pupil size, this was not notably reflected in differences in power or intertrial coherence. This suggests that the decoder relied on more subtle differences in the beta band, such as modulation of ERPs that are expressed in this frequency range (also see <u>EEG: lateralized ERPs</u>).

**Figure 7.**
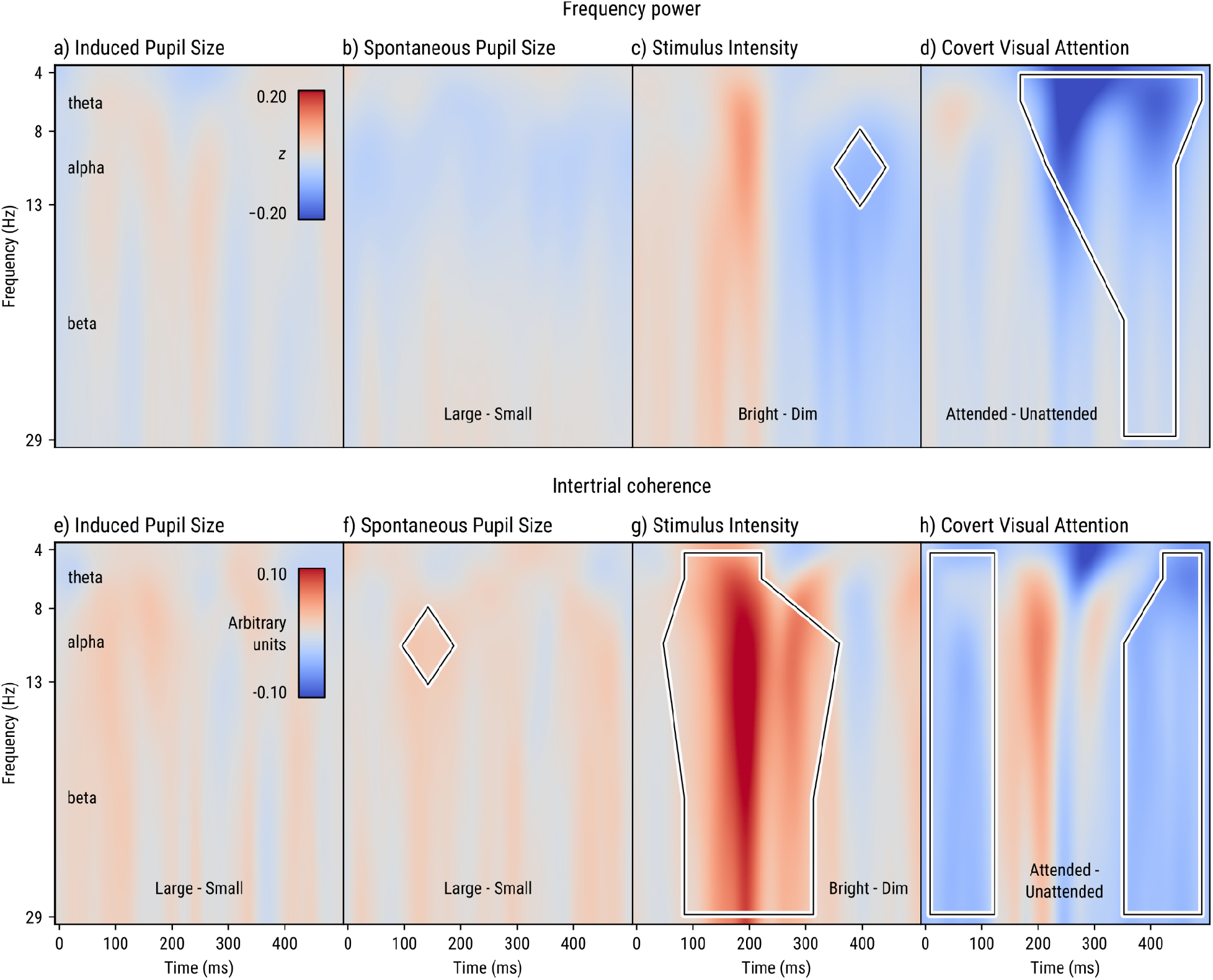
Time-frequency plots as a function of time since target onset (x-axis) and frequency (y-axis) for power as a function of a) induced pupil size, b) spontaneous pupil size, c) stimulus intensity, and d) covert visual attention; and intertrial coherence as a function of e) induced pupil size, f) spontaneous pupil size, g) stimulus intensity, and h) covert visual attention. Positive values are indicated in red; negative values are indicated in blue. Outlines provide a rough indication of frequencies and times where significant contrasts are observed (see main text for details).

#### Spontaneous differences in pupil size weakly affect coherence (Fig. 7b,f)

Although the decoding results showed that beta frequencies, mainly around 22 Hz, were somewhat affected by spontaneous pupil size, this was again not notably reflected in differences in power or intertrial coherence, again suggesting that the decoder relied on other effects expressed in this frequency range. However, there was a slight increase in alpha intertrial coherence for spontaneously large as compared to small pupils (*p* = .018), which matched the slight reliance on alpha frequencies as revealed in the decoding results.

#### Stimulus intensity affects alpha power and coherence in all frequency bands (Fig. 7c,g)

Decoding results showed that alpha frequencies, mainly around 10 Hz, were strongly affected by stimulus intensity. This was reflected in an decrease in alpha power for bright as compared to dim trials with a peak around 400 ms (*p* < .001). This was also strongly reflected in intertrial coherence, which was larger for bright as compared to dim trials in all frequency bands (all *p* < .001), with a peak around 200 ms in the alpha band.

#### Covert visual attention strongly affects power and coherence mainly in theta but also in alpha and beta frequency bands (Fig. 7d,h)

Decoding results showed that theta frequencies, mainly around 5 Hz, were somewhat affected by covert visual attention. This was also strongly reflected in power, which was smaller for attended as compared to unattended trials for most of the time window and in all frequency bands (theta and alpha: *p* < .001; beta: *p* = .006), with a peak around 250 ms in the theta band. This was also reflected in intertrial coherence, although here the pattern was more complex, characterized by an initial very rapid decrease in coherence for attended as compared to unattended trials (all bands: *p* < .001), which disappeared (and numerically even reversed, though this was not reliable) around 200 ms, followed by a continuation of the decrease (theta: *p* = .014; alpha: *p* = .001; beta: *p* < .001), with a trough around 400 ms in the alpha and beta bands.

### Pupil constriction

We investigated how the strength of pupil constriction in response to the target was affected by our four factors (Fig. 8). To test this, we used cross-validation to localize the sample at which each main effect was strongest within the 500 - 700 ms post-target window, and then conducted a single linear mixed-effects analysis (see `time_series_test.lmer_crossvalidation_test`)^7^. Pupil size was baseline corrected relative to a 50 ms pre-target baseline period. Induced pupil size, spontaneous pupil size, stimulus intensity, and covert visual attention were included as fixed effects; no interactions were included; by-participant random intercepts and slopes were included. Since pupil responses are typically slower than responses as measured through EEG, we used a slightly longer window from 0 to 1000 ms after target onset.

**Figure 8.**
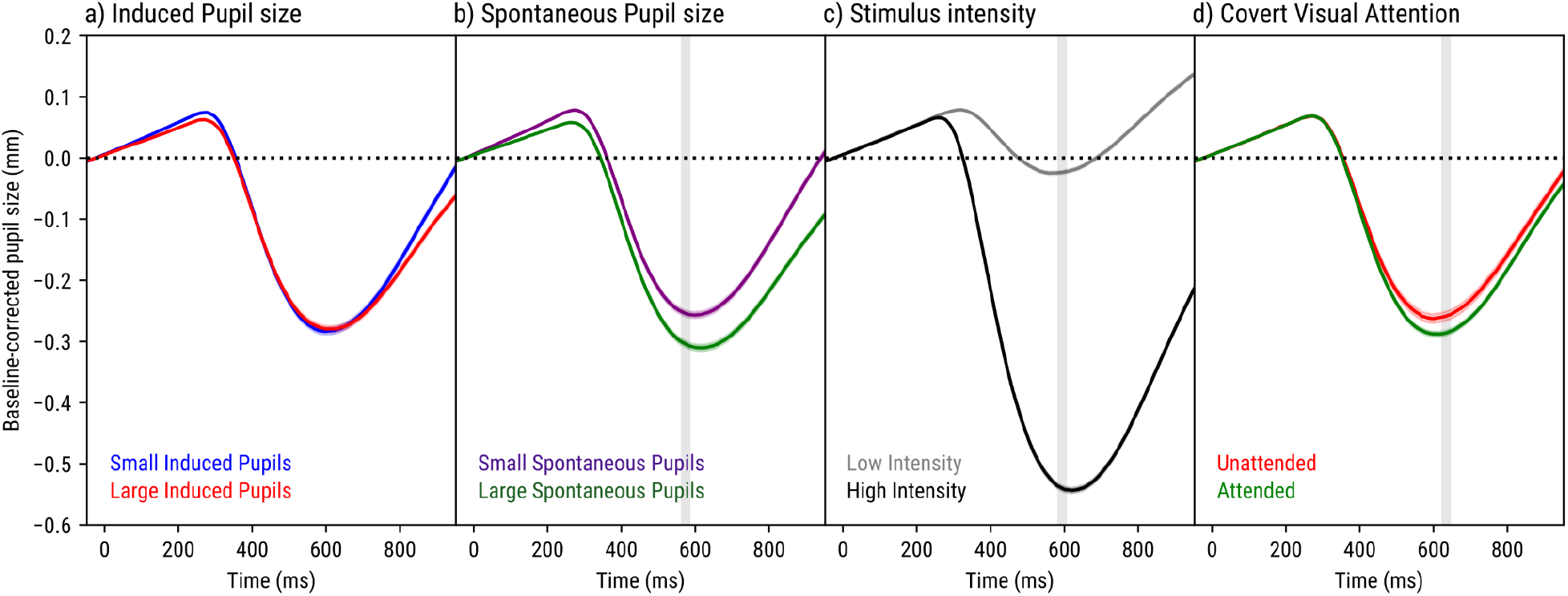
Target-evoked pupil responses as a function of time (x-axis) and a) induced pupil size, b) spontaneous pupil size, c) stimulus intensity, and d) covert visual attention. Responses are baseline corrected, and therefore induced and spontaneous differences in baseline pupil size are not visible here. Vertical bands indicate peak differences (see main text for details).

#### Induced differences in baseline pupil size do *not* affect pupil constriction (Fig. 8a)

Pupil constriction was equally strong for induced large and small pupils (*z* = 1.37, *p* = .170); that is, even though the inducers affected the absolute size of the pupil, they did not affect how much the size of the pupil changed in response to the target. Since approximately 20% more light entered the eye on induced large as compared to induced small pupil trials (see <u>Supplementary results</u>), this is not a trivial result, and suggests a form of brightness constancy that already operates on the level of the pupil light response.

#### Spontaneous differences in baseline pupil size do affect pupil constriction (Fig. 8b)

Pupils constricted more strongly for spontaneously large pupils as compared to small pupils (*z* = −6.99, *p* < .001). Given that this effect did not occur for induced differences in baseline pupil size, it likely did not result from increased light influx (which was about 23% for spontaneous large vs small pupils, comparable to the 18% for induced large vs small pupils), but rather was mediated by increased arousal; that is, large baseline pupil sizes reflect increased attentiveness, in turn resulting in stronger pupil constriction, analogous to the effect of covert visual attention described below.

#### Stimulus intensity affects pupil constriction (Fig. 8c)

Pupils constricted more strongly to high-intensity targets as compared to low-intensity targets (*z* = −18.68, *p* < .001). This reflects the typical pupil light response.

#### Covert visual attention affects pupil constriction (Fig. 8d)

Pupils constricted more strongly to targets presented at attended locations as compared to unattended locations (*z* = −5.52, *p* < .001). This reflects the modulation of the pupil light response by covert visual attention (Binda & Murray, 2015).

## General discussion

Here we report that pupil size causally affects visual processing as measured through EEG. We experimentally manipulated pupil size by presenting either an equiluminant blue or red inducer prior to each trial (Wardhani et al., 2022); prolonged exposure to blue light activates ipRGCs, which in turn triggers sustained pupil constriction (Gamlin et al., 2007; Mathôt, 2018), thus providing an unobtrusive way to manipulate pupil size. We compared the effects of both induced and spontaneous differences in pupil size to the effects of stimulus intensity and covert visual attention, all of which have been reported to enhance visual processing as measured by increased detectability of stimuli (Carrasco et al., 2000) and increased SSVEPs (Müller et al., 1998; Suzuki et al., 2019).

We applied neural-network decoding of EEG signals following target onset to quantify whether and how visual processing was affected by induced pupil size, spontaneous pupil size, stimulus intensity, and covert visual attention (Schirrmeister et al., 2017). We found that all of these factors could be decoded highly reliably, with occipital and parietal electrodes carrying most information. However, although the topographical distribution was roughly similar for all four factors, there were also pronounced differences, mainly in terms of which frequency bands carried most information: decoding of induced pupil size mainly relied on beta frequencies (22 Hz peak); spontaneous pupil size also mainly relied on beta frequencies (22 Hz peak) and to a lesser extent on alpha frequencies; stimulus intensity mainly relied on alpha frequencies (10 Hz peak) and to a lesser extent on theta frequencies; and covert visual attention mainly relied on theta frequencies (5 Hz peak) and to a lesser extent also on alpha frequencies. These dissociations, corroborated by qualitative differences in lateralized ERPs, stimulus-evoked pupil constrictions, and a lack of cross-decoding between factors, suggest that induced pupil size, spontaneous pupil size, stimulus intensity, and covert visual attention all have qualitatively different effects on visual processing.

A first important implication of our results is that pupil size causally affects visual processing as measured through EEG, especially in the beta band. This is reminiscent of correlational studies that observed a positive correlation between spontaneous pupil-size changes and beta power (Keegan & Merritt, 1995; Podvalny et al., 2021; Waschke et al., 2019), and extends these findings by suggesting that this link is causal. One important difference between our study and these previous studies is that previous studies all focused on activity during rest or a pre-stimulus baseline, rather than on activity following target onset; another important difference is that, although we found a highly pronounced reliance on beta activity for decoding, we did not find differences in overall beta power or intertrial coherence; this suggests that pupil size affected beta-band activity in more subtle ways, such as by modulating ERPs that are expressed in this frequency range, and which our ERP analyses also highlighted as being affected by pupil size. However, decoding and time-frequency analyses are fundamentally different, which makes it difficult to directly compare them. For example, decoding is done for each participant separately, which means that it is able to pick up on idiosyncratic effects that are systematic within participants but that differ across participants; in contrast, time-frequency analysis is done across participants, which means that it is only able to pick up effects that are systematic across participants. Furthermore, decoding and time-frequency analyses require different preprocessing steps. Nevertheless, although there are still many details to be worked out, the picture that emerges from our results and others is that pupil size causally affects activity in the beta band—so what might this reflect, in terms of underlying cognitive processes?

One dominant view relates beta activity to motor activity, typically such that increased beta power is associated with increased tonic muscle contractions (Engel & Fries, 2010; Jenkinson & Brown, 2011); this raises the possibility that the link between pupil size and beta activity reflects oculomotor control of the iris muscles, such as an increased tonic contraction of the iris dilator muscle during pupil dilation. Another—not mutually exclusive—view relates beta activity to visual perception such that increased phase coherence (but not power) is associated with enhanced visual perception (Bressler & Richter, 2015; Hanslmayr et al., 2007); this second view raises the possibility that the link between pupil size and beta activity reflects facilitation of visual perception as a result of the increased light influx that accompanies increased pupil size.

These are testable hypotheses for future studies in which, for example, pupil size and stimulus detectability could be dissociated in order to test which of the two is most strongly related to beta activity; if pupil size affects beta activity more strongly than stimulus detectability does, then the effect on beta is likely related to motor control, whereas the opposite pattern would suggest that the effect is more closely related to perception. A related important aim for future research will be to better characterize these effects on beta activity, and to better understand how induced effects, such as modulations of power or intertrial coherence, which previous studies have mainly focused on (Keegan & Merritt, 1995; Podvalny et al., 2021; Waschke et al., 2019), relate to evoked effects, such as modulations of ERPs expressed in a particular frequency range, which our results seem to point towards.

A second important implication of our results is that there is a correlation between intertrial coherence in the alpha band and spontaneous pupil size. This correlation is weak and should be replicated, especially given the exploratory nature of our study. However, assuming that it is reliable, it is likely mediated by arousal (or increased attentiveness) rather than by pupil size per se, because it was not observed for induced pupil size, which presumably does not affect arousal. This correlation is likely analogous to (yet far weaker than) the effect of covert visual attention on alpha power and intertrial coherence (see also Cómez et al., 1998): the more a stimulus is attended, either because it has been cued (covert visual attention) or because the participant just happens to be in a state of increased vigilance (spontaneous pupil size), the more alpha intertrial coherence increases.

Although we did not observe a reliable correlation between alpha power and spontaneous pupil size, power and intertrial coherence are related concepts (van Diepen & Mazaheri, 2018). Therefore, it is tempting to link the observed correlation between alpha intertrial-coherence and spontaneous pupil size to the well-established negative correlation between pre-stimulus (as opposed to post-stimulus) alpha power and visual-detection performance (Ergenoglu et al., 2004; Thut et al., 2006); however, this link is complicated by recent reports of both positive (Ceh et al., 2020; Montefusco-Siegmund et al., 2022) and negative (Hong et al., 2014) correlations between pre-stimulus alpha power and spontaneous pupil size. In our view, pre-stimulus alpha power and spontaneous pupil size likely reflect different dimensions of attention; specifically, pre-stimulus alpha seems suppressed when attention is directed to external visual stimuli (Kraus et al., 2023; Peylo et al., 2021; van den Berg et al., 2014), as during a spatial cueing task, and enhanced when attention is directed to internal states, as during mind wandering (Ceh et al., 2020; Ray & Cole, 1985); in contrast, pupil size seems to reflect the “intensity” of attention regardless of whether this is directed internally or externally (Just & Carpenter, 1993; Kraus et al., 2023; Mathôt, 2018). Therefore, whether pre-stimulus alpha power and spontaneous pupil size correlate negatively or positively with each other, or not at all, likely depends on the context of the task. An important aim for future research will be to better understand whether and how pupil size, pre-stimulus alpha (not addressed in the analyses presented here), post-stimulus alpha, and behavioral performance correlate, and to what extent these correlations are mediated directly by pupil size or indirectly by arousal.

A third important implication of our results is that pupil size should not be considered as just a marker of cognitive state without also considering that pupil size itself affects visual processing. This is particularly relevant to the growing literature on ‘pupil-linked arousal’, which is built on the finding that spontaneous fluctuations in pupil size reflect—among other things and other brain areas (Joshi et al., 2016; Megemont et al., 2022)—activity in the locus-coeruleus-norepinephrine (LC-NE) system (e.g., Alnaes et al., 2014; Aston-Jones & Cohen, 2005; de Gee et al., 2014; Hong et al., 2014; Murphy et al., 2011, 2014; Urai et al., 2017; Waschke et al., 2019). Our findings suggest that researchers should be especially careful when interpreting correlations between spontaneous fluctuations in pupil size and performance on a visual task in terms of LC-NE activity.

Finally, we have so far assumed that our pupil-size induction procedure is a pure manipulation of pupil size, such that there is one causal arrow from the red/ blue inducers to pupil size, and a second causal arrow from pupil size to visual processing; this assumption is the justification for our use of the word ‘causal’ in the title of the current manuscript. By looking at the effects of both induced and spontaneous changes in pupil size, and by contrasting these with the well-known effects of stimulus intensity and covert visual attention, we have made an important first step towards understanding the causal role of pupil size in visual processing. However, future studies are necessary to further establish the validity of this procedure. For example, our pupil-size induction procedure could be combined with concurrent heart-rate and skin-conductance measurements to confirm that the red/ blue inducers do not differentially affect arousal or fatigue, which could in turn affect visual processing in ways that are unrelated to pupil size. Future studies could also use different ways to manipulate pupil size; for example, eye drops (tropicamide) are a minimally invasive way to pharmacologically dilate the pupil; however, eye drops also affect lens accommodation, which means that they are only suitable for experiments that can be done with blurry vision. In general, in order to fully understand how pupil size causally affects visual processing, future studies will need to develop and use a range of different, well-validated techniques to manipulate pupil size, one of which will be the red/ blue induction procedure that we have used here.

In sum, we found that pupil size affects visual processing as measured through EEG, mainly over occipital and parietal electrodes, and mainly in beta frequencies; we observed this for both induced and—although less clearly—spontaneous pupil-size changes, which suggests that this reflects a direct causal effect of pupil size on visual processing. We further found that spontaneous (but not induced) pupil size is negatively correlated with power and positively with intertrial coherence in the alpha band; these correlations are likely mediated by vigilance/ arousal (Loewenfeld, 1958). Finally, we found that the effect of pupil size on visual processing is qualitatively different from the effects of stimulus intensity and covert visual attention, and is therefore a unique factor that shapes early visual processing.

## Open-practices statement

Experimental data, the experiment script, and analysis scripts can be found at https://osf.io/gjpkv/

## Acknowledgements

This research was supported by the Innovational Research Incentives Scheme VIDI (VI.Vidi.191.045) from the Dutch Research Council (NWO).

## Supplementary results

### Red and blue inducers were successfully matched

At the end of the calibration procedure, pupil responses to red (*xyY* color coordinates: 0.15, 0.07, 6.74 cd/m^2^) and blue (*xyY*: 0.65, 0.34, 9.84 cd/m^2^) inducers were equally strong; that is, the calibration procedure had successfully matched the intensity of the red and blue inducers as reflected by the strength of the initial (but not the sustained) pupil constriction (Fig. 9). To test this, we took mean pupil size during the 0.95 - 1.05 s window for the last trial for each color and participant separately. Next, we conducted a default Bayesian paired-samples t-test, which revealed substantial evidence for the null hypothesis (BF_01_ = 3.57).

**Figure 9.**
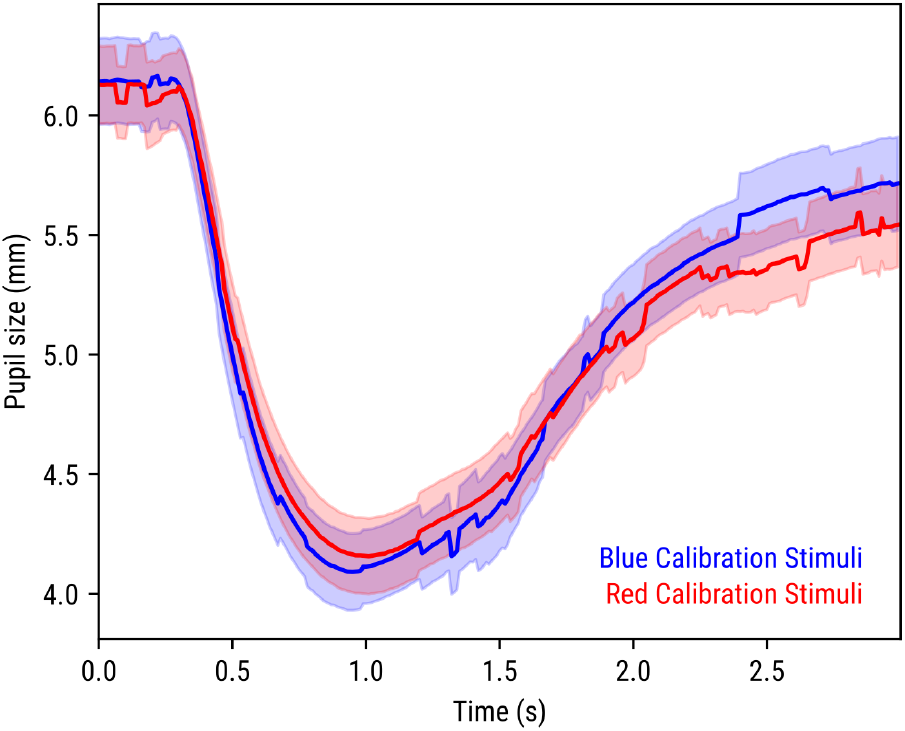
Pupil responses to brief (1 s) exposure to red/ blue stimuli during calibration. Lines are averages of the last trials of each color for each participant. Error bands reflect standard errors.

### Behavioral performance was affected by cue validity, but not by inducer color

We analyzed behavioral performance to verify that a) participants indeed directed their attention towards the cued side (an effect of covert visual attention), and b) the staircase procedure successfully kept performance constant between validly cued trials of red- and blue-inducer blocks (no effect of induced pupil size; see Fig. 10). To do so, we conducted a logistic linear mixed effects analysis with response accuracy (0, 1) as dependent measure, induced pupil size and covert visual attention and the induced pupil size × covert visual attention interaction as fixed effects, and random by-participant intercepts and slopes for all fixed effects; this revealed a main effect of covert visual attention (*z* = 2.747, *p* = .006), such that accuracy was higher on attended trials than on unattended trials, but no main effect of induced pupil size (*z* = 0.314, *p* = .754), and no interaction between induced pupil size and covert visual attention (*z* = 0.422, *p* = .673). We also conducted an analogous (non-logistic) analysis using response time (for correct trials only) as dependent measure; this again revealed a main effect of covert visual attention (*t* = −7.714, *p* < .001), such that responses were faster on attended cued trials than on unattended cued trials, but no main effect of induced pupil size (*t* = −0.181, *p* = .858), and no interaction between induced pupil size and covert visual attention (*t* = −0.106, *p* = .916).

**Figure 10.**
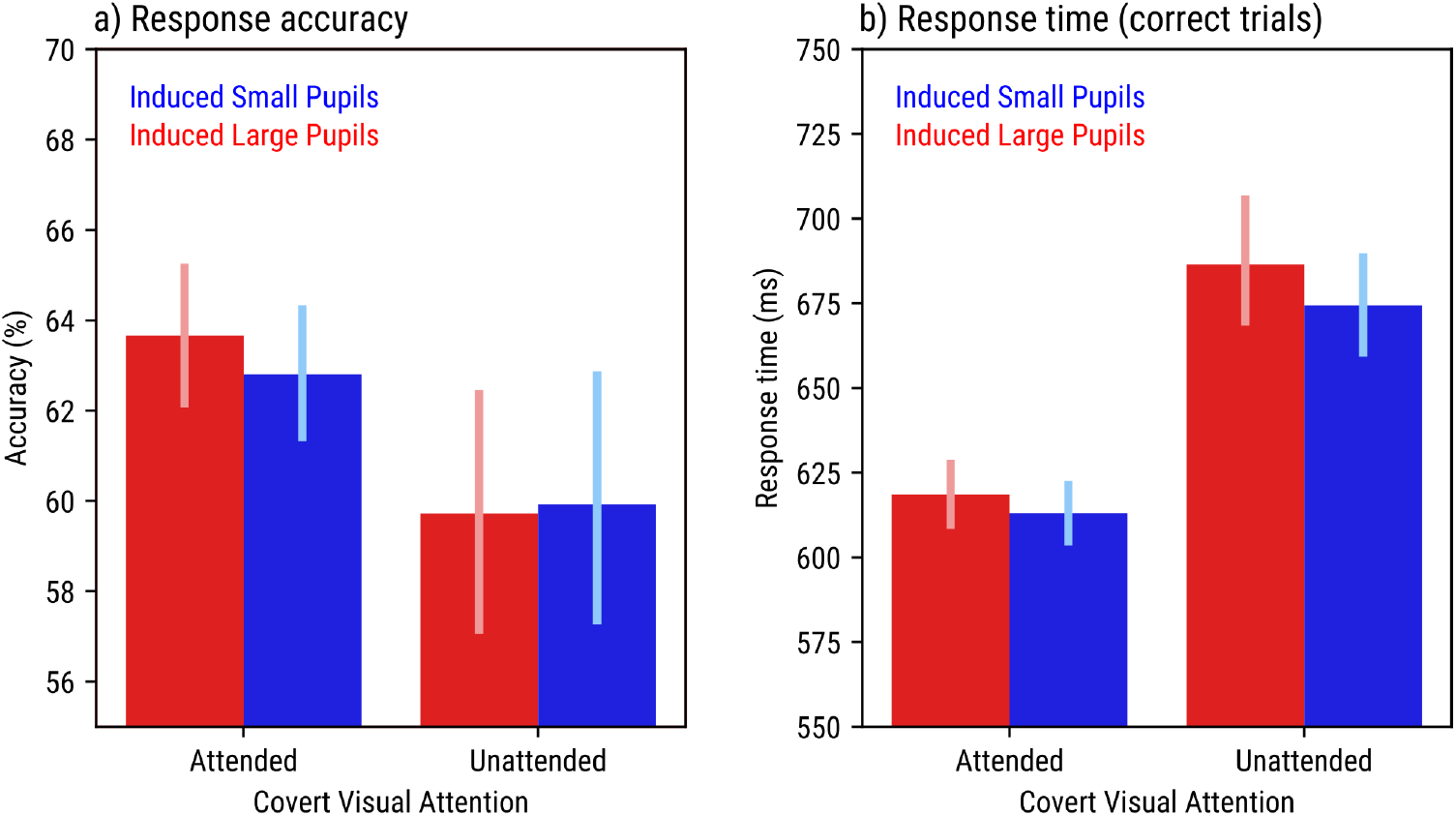
Behavioral response accuracy (a) and response time (b) as a function of induced pupil size and covert visual attention. Error bars represent 95% bootstrap confidence intervals.

### Microsaccade rate correlates with spontaneous pupil size, but not with other factors

Microsaccades are small saccadic eye movements (Dimigen et al., 2009; Yuval-Greenberg et al., 2008). In experiments such as the current one, the presentation of a stimulus is typically followed by a period of about 200 ms during which there are very few microsaccades, followed by a pronounced increase in microsaccade rate. Because of this predictable change in microsaccade rate following stimulus onset, microsaccades may evoke EEG responses that seem to be locked to the stimulus while they are in fact locked to microsaccades (Yuval-Greenberg et al., 2008). Microsaccades are also correlated with pupil size (Wang & Munoz, 2021). Taken together, this means that all of the results presented in the main text could in principle be mediated by differences in microsaccade rate between the various conditions.

To test the contribution of microsaccades to the results, we investigated how microsaccade rate after target onset was affected by our four factors (Fig. 11). We determined the rate of saccades smaller than 2° using `pycrosaccade`^8^, a Python package that implements a well-established algorithm for detecting microsaccades (Engbert & Kliegl, 2003). We then conducted a cross-validation test as described above for pupil constriction, with the only difference that we now analyzed the full 1000 ms following target onset (see Fig 11).

**Figure 11.**
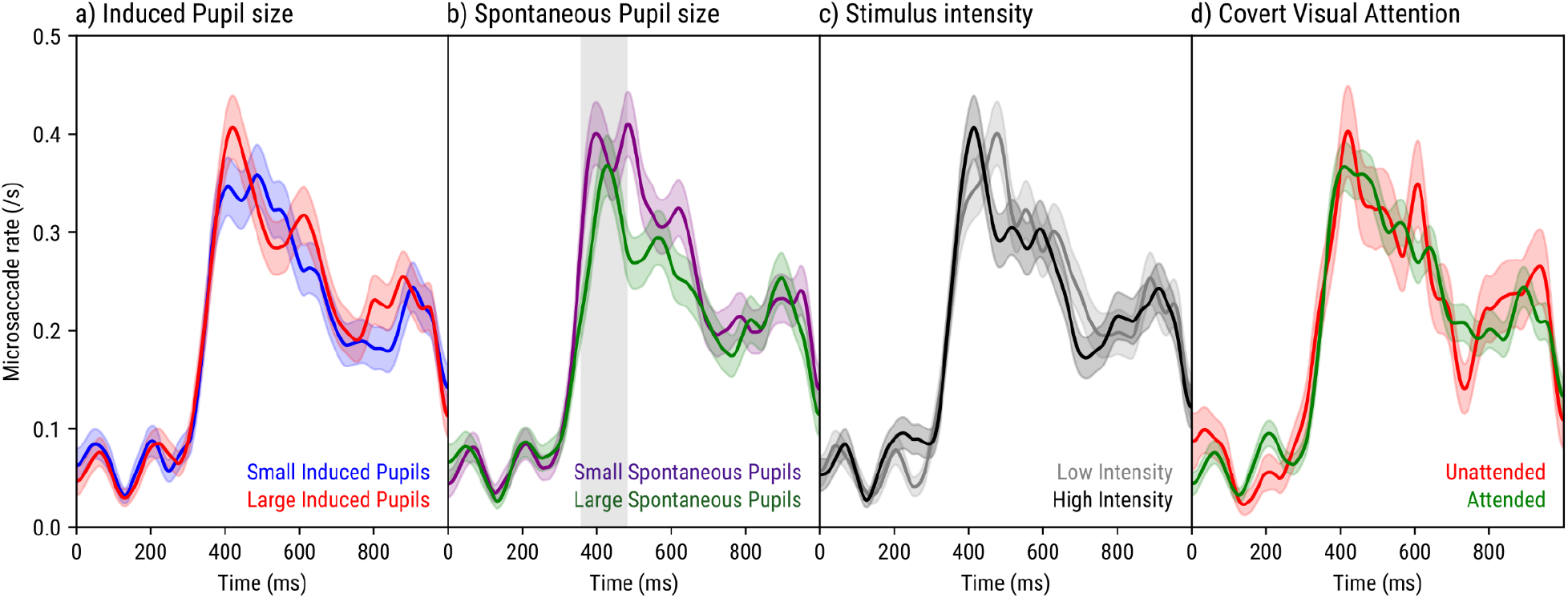
Microsaccade rate as a function of time (x-axis) and a) induced pupil size, b) spontaneous pupil size, c) stimulus intensity, and d) covert visual attention. The vertical shaded area indicates a maximum difference between conditions in roughly that period.

Microsaccade rate was very low immediately after stimulus onset, and increased sharply after about 300 ms; this pattern is similar, though somewhat slower and with lower overall rates, to what has previously been reported for post-stimulus microsaccades (Yuval-Greenberg et al., 2008). We did not observe pronounced differences in microsaccade rate between most conditions, except for a slight difference such that microsaccade rate was lower for large as compared to small spontaneous pupils (*z* = −1.97, *p* = .049). However, this effect should be interpreted with caution given that it was weak and part of an exploratory analysis.

Importantly, the fact that microsaccade rate was not notably affected by our factors, with the possible exception of spontaneous pupil size, suggests that our main results are not mediated by microsaccade rate.

### Non-linear effects of spontaneous pupil size on lateralized ERPs

For the main analyses, we distinguished only between small and large spontaneous pupil size, thus implicitly making the assumption that the relationship between spontaneous fluctuations in pupil size and EEG activity should be approximately linear. However, previous work has found that spontaneous pupil size often shows an inverted-U-shaped relationship to (EEG-correlates of) task performance, such that best performance is observed with fairly large pupils, whereas both *very* large pupils and small pupils are associated with poor performance (e.g., Brink et al., 2016; Waschke et al., 2019). This finding is generally interpreted in terms of the Yerkes-Dodson law (Yerkes & Dodson, 1908), which similarly posits that best performance is observed for fairly high levels of arousal, whereas both very high levels of arousal and low levels of arousal are associated with poor performance.

Therefore, we re-analyzed lateralized ERPs as a function of spontaneous pupil size (similar to Fig. 6b), but this time for visualization using five bins rather than two in order to reveal any non-linearities in the relationship between spontaneous pupil size and lateralized ERPs (Fig. 12a). We zoomed in on the 250-300 ms window that we had previously found to be affected by spontaneous pupil size (Fig. 12b). Qualitatively, we do indeed observe an inverted-U-shaped relationship such that lateralized ERPs are least pronounced for intermediate-to-large pupils (around 5.5 mm) and most pronounced for small and very large pupils.

**Figure 12.**
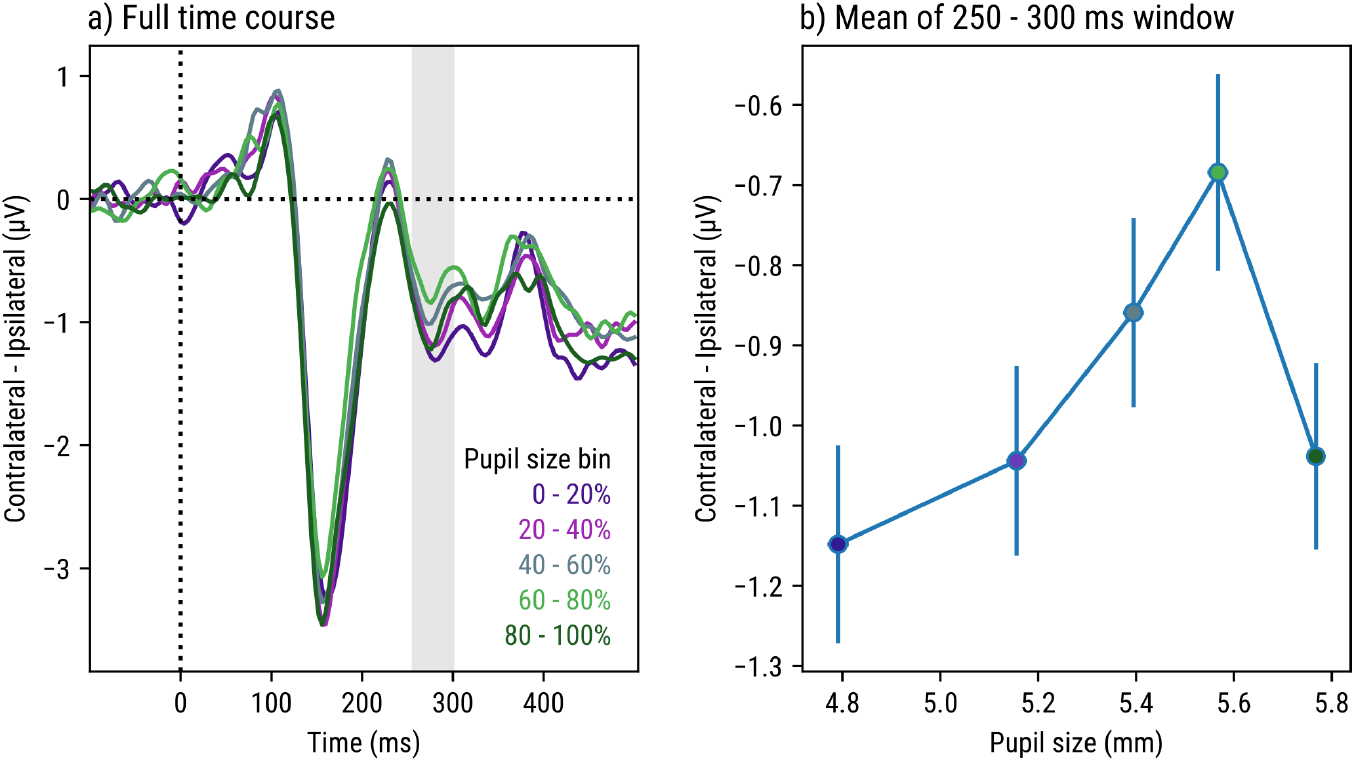
a) Lateralized target-evoked ERPs (baseline corrected) as a function of time (x-axis) and spontaneous pupil size, split into five bins. b) The mean voltage difference between contralateral and ipsilateral electrodes during the 250 - 300 ms window (indicated in a as a vertical shading) as a function of spontaneous pupil size. Error bars represent standard errors.

We then conducted a linear-mixed-effects analysis using the mean voltage of contralateral minus ipsilateral electrodes during the 250 - 300 ms windows as the dependent variable, and linear spontaneous pupil size (z-scored per participant) and quadratic spontaneous pupil size (i.e. z-scored spontaneous pupil size raised to the power of two) as fixed effects, and by-participant random intercepts. This revealed an effect of linear spontaneous pupil size (*z* = 2.60, *p* = .009) but no effect of quadratic spontaneous pupil size (*z* = 0.66, *p* = .507).

Taken together, the relationship between spontaneous pupil size and lateralized ERPs qualitatively resembled a non-linear inverted-U shape. However, statistical testing revealed that the non-linear (quadratic) component of this relationship (unlike the linear component) was not reliable.

### Decoding of induced pupil size does not rely on proximity in time

A potential issue when decoding induced pupil size is that this factor is varied between blocks, rather than varied randomly from trial to trial. This means that a trial with large induced pupils is closer in time to other trials with large induced pupils than to other trials with small induced pupils. In turn, this means that the decoder may learn to pick up on slow fluctuations in the EEG signal that do not have anything to do with pupil size per se, but that due to proximity in time nevertheless allow for above-chance decoding of induced pupil size.

To account for this, we ran a control analysis in which we split each block into two halves. We then trained on the first halves and tested on the second halves while excluding the very last half altogether. If we use L to represent large-induced-pupil block halves and S to represent small-induced-pupil block halves, and given a participant who started with a large-induced-pupil block, this training-and-testing scheme looks as follows:

`Train: L-S-L-S-`

`Test: -L-S-L--`

This scheme has the property that each test session is flanked by both an L and an S block, which prevents the decoder from relying on proximity in time. We complemented this with the following scheme, which has the same property:

`Train: -L-S-L-S`

`Test: --S-L-S-`

For each participant separately, we used both of these schemes to decode induced pupil size, and then took the average of the decoding accuracy of the two schemes. Despite this not being a very powerful decoding procedure as compared to traditional four-fold cross-validation, induced pupil size was nevertheless decoded far above chance (62%; one-sample t-test against chance level: *t* = 4.27, *p* < .001).

We also conducted yet another control analysis in which we split the entire experimental session into two halves, and trained and tested on those:

`Train: LLSS----`

`Test: LLSS`

And:

`Train: LLSS`

`Test: LLSS----`

Even when using this highly suboptimal training-and-testing scheme, in which the testing data is far apart in time from the training data, we were able to decode induced pupil size above chance (57%, *t* = 2.41, *p* = 0.023).

## Footnotes

1 A far larger number of studies have looked at correlations, often in the context of “pupil-linked arousal”. We will return to this in the <u>General discussion</u>.

2 https://github.com/smathot/eeg_eyetracking_parser

3 https://braindecode.org/

4 As described under <u>Participants</u>, two additional participants were tested and replaced because they did not show this effect. It is currently unclear whether this reflects measurement noise or systematic individual differences in the extent to which people are affected by red and blue inducers. However, overall the effect is highly systematic.

5 The analysis source code contains sanity-check analyses showing that the cross-decoding analysis is able to pick up similarities between factors that should be very similar, such as by training on the effect of Covert Visual Attention on Low Intensity trials, and then decoding the effect of Covert Visual Attention on High Intensity trials.

6 The analysis source code again contains additional analyses focusing on different sets of electrodes and different contrasts.

7 Since the pupil response in this case consists of only a single component (a pupil light response), there is no need to conduct a cluster-based permutation test, which is extremely computationally intensive when combined with linear-mixed effects modeling (see Mathôt & Vilotijević, 2022).

8 https://github.com/robbertmijn/pycrosaccade

